# Microfluidic Platform for Drug Response Profiling in NSCLC Patient-Derived Organoids

**DOI:** 10.64898/2026.06.17.733025

**Authors:** Qiyue Luan, Alireza Rahnama, Ines Pulido, Mario Raspini, Jian Zhou, Takeshi Shimamura, Ian Papautsky

**Affiliations:** Department of Biomedical Engineering, University of Illinois Chicago, Chicago, IL, USA; Department of Surgery, University of Illinois Chicago, Chicago, IL, USA; University of Illinois Cancer Center, Chicago, IL, USA; Department of Cardiovascular and Thoracic Surgery, Rush University Medical Center, Chicago, IL, USA; Department of Anatomy and Cell Biology, Rush University Medical Center, Chicago, IL, USA

## Abstract

Tumor models that recapitulate 3D architecture are essential for understanding how cellular organization and microenvironmental interactions govern therapeutic response in human cancers. Here, we developed a microfluidic microphysiological system that enables controlled and scalable culture and drug testing of non-small cell lung cancer spheroids and patient-derived organoids. The platform integrated U-shaped microwells with dual-channel loading to support de novo spheroid formation, efficient trapping of pre-formed spheroids, and loading of intact organoids with reduced size heterogeneity. Tumor spheroids and organoids maintained high viability and structural integrity during long-term on-chip culture, and constrained microscale confinement produced ellipsoidal geometries that deviate from idealized spherical assumptions. Baseline genotype-dependent responses to KRAS G12C and EGFR inhibitors were preserved across agarose and microfluidic formats, establishing a validated reference state. Building on this baseline, fibroblast- and endothelial-derived cues consistently attenuated responses to targeted therapies across conditioned media, mixed co-culture, and spatially organized configurations. Resistance phenotypes converged on a dominant role for paracrine signaling, while increasing architectural complexity primarily enhanced morphological fidelity rather than altering therapeutic response. These findings establish a microphysiological framework that decouples tumor-intrinsic drug sensitivity from microenvironment-mediated modulation, enabling the systematic evaluation of paracrine resistance mechanisms in NSCLC.

## 1. Introduction

The tumor microenvironment (TME) plays a critical role in shaping therapeutic response and driving resistance in non-small cell lung cancer (NSCLC).^1–9^ Stromal components, including fibroblasts and endothelial cells, secrete growth factors, chemokines, and cytokines that regulate tumor survival, proliferation, and sensitivity to targeted therapies. These paracrine interactions can limit drug penetration, activate compensatory signaling pathways, and promote phenotypic persistence. However, conventional preclinical models such as 2D monolayers and murine xenografts fail to recapitulate the spatial organization, diffusional constraints, or dynamic tumor-stromal communication observed in human tumors, limiting their translational relevance.^10,11^ Although *in vivo* models partially capture tumor complexity, they are costly and poorly scalable, particularly for patient-specific studies. As a result, there remains a need for experimentally tractable platforms that preserve human tumor heterogeneity while reducing material requirements and experimental burden.

Patient-derived organoids (PDOs) provide physiologically relevant models by preserving key genomic and phenotypic features of the parental tumor.^12,13^ Despite this advance, their application in mechanistic drug studies is constrained by technical limitations. Standard Matrigel dome cultures generate PDOs with substantial size heterogeneity, introducing variability in drug uptake, diffusion, and viability measurements, and complicating quantitative comparisons.^14–16^ In addition, existing co-culture strategies typically combine PDOs with stromal cells in an unstructured formats, limiting the ability to distinguish paracrine signaling from contact-dependent interactions and to systematically assess how increasing architectural complexity influences drug response.

To address these limitations, we developed a microfluidic microphysiological system (MPS) that enables quantitative analysis of drug responses in oncogene-driven NSCLC under controlled microenvironmental conditions. The system integrates U-shaped microwells that support formation of tumor spheroids and culture of size-controlled PDOs, reducing heterogeneity and enabling robust statistical comparisons.^17,18^ Importantly, the platform preserves baseline tumor-intrinsic drug sensitivity while allowing controlled introduction of fibroblast- and endothelial-derived cues, decoupling intrinsic therapeutic response from microenvironment-mediated modulation. This design provides a systematic framework for isolating soluble microenvironmental factors that influence the efficacy of targeted agents, including KRAS and EGFR inhibitors. The system further supports multiple TME configurations, including exposure to conditioned media, mixed tumor-stromal aggregates, and spatially organized microfluidic co-cultures with defined compartmentalization. These formats enable interrogation of TME signaling across increasing levels of architectural complexity while maintaining control over tumor size, positioning, and drug delivery. In addition, the platform is compatible with on-chip imaging, immunofluorescence, and recovery of organoid material, supporting flexible mechanistic and downstream molecular analyses.

Using this platform, we investigated how fibroblast and endothelial-derived cues influence responses to targeted therapies in NSCLC cell line spheroids and PDOs. By first establishing baseline tumor intrinsic drug sensitivity and then introducing defined microenvironmental perturbations, we assessed resistance phenotypes across conditioned media, mixed co-culture, and spatially organized microphysiological configurations. Drug response patterns observed on-chip were consistent with those measured using parallel agarose microwell systems, supporting the platform’s robustness. Together, these studies establish a controlled experimental framework for distinguishing TME-driven resistance mechanisms from effects arising from increasing tissue architectural complexity. Accordingly, this study aimed to 1) validate a microfluidic platform that preserves baseline tumor-intrinsic drug sensitivity, 2) systematically isolate paracrine-mediated resistance using stromal and endothelial cues, and 3) evaluate how increasing architectural complexity influences resistance phenotypes.

## 2. Materials and Methods

### 2.1. Materials

Cell lines A549 (KRAS G12V mutant), NCI-H358 (KRAS G12C mutant, hereafter referred to as H358, NCI-H1975 (EGFR L858R/T790M mutant, hereafter referred to as H1975), and WI-38 (lung fibroblast) were acquired from ATCC (Manassas, VA, USA). Human Umbilical Vein Endothelial Cells (HUVECs) were acquired from Lonza (Basel, Switzerland).

All cell lines were maintained according to the suppliers’ recommended protocols. NSCLC PDOs F231 and F671 were obtained from the NCI Patient-Derived Models Repository (PDMR). RPMI 1640 medium with L-glutamine, EMEM, DMEM, trypsin–EDTA (0.25%), 1× phosphate-buffered saline (PBS), Dispase II, and 24/48-well cell culture plates were purchased from Fisher Scientific (Waltham, MA, USA). Fetal bovine serum (FBS) was purchased from GeminiBio Inc. (West Sacramento, CA, USA). Antibiotic-antimycotic solution (100×) and Calcein AM were purchased from Invitrogen (Carlsbad, CA, USA). Agarose powder was sourced from Sigma-Aldrich (St. Louis, MO, USA), and propidium iodide from Alfa Aesar (Haverhill, MA, USA).

Hoechst 33342 solution was purchased from Thermo Fisher Scientific (Waltham, MA, USA). The targeted inhibitors adagrasib and osimertinib were purchased from MedChemExpress (Monmouth Junction, NJ, USA). The CellTiter-Glo 3D Cell Viability Assay was obtained from Promega (Madison, WI, USA). Basement Membrane Extract Type 2 (BME2) was purchased from R&D Systems (Minneapolis, MN, USA). Sylgard 184 polydimethylsiloxane was purchased from Ellsworth (Germantown, WI, USA). SUEX negative photoresist dry films were purchased from DJ Microlaminates (Sudbury, MA, USA).

### 2.2. Microfluidic device fabrication

The microfluidic device consisted of three PDMS layers. The bottom layer contained U-shaped microwells for capturing spheroids or organoids. The middle layer contained flow channels with integrated circular culture chambers, and the top layer contained reservoirs to supply media to the device inlets. Microwells in the bottom layer had a diameter of 400 µm, a depth of approx. 250 µm, and were arranged in parallel rows with a center-to-center spacing of 600 µm. These dimensions were selected to accommodate intact NSCLC spheroids and PDOs while limiting vertical displacement during handling and perfusion.

The bottom layer was fabricated using polycarbonate (PC) masters to form U-shaped microwells, following the previously reported process.^19^ Briefly, PDMS replicas were first produced from a 350 µm thick SUEX dry photoresist mold fabricated as described previously^20^ to form flat-bottom microwells. Briefly, SUEX films were laminated onto 3 in silicon wafers, photolithographically patterned, and hard-baked at 200°C for 2 h. PDMS was prepared at a 10:1 (v/v) monomer-to-curing-agent ratio, degassed, cast onto the master molds, and cured at 80°C for 2 h. These PDMS replicas were then used to shape PC slabs. PC slabs were cleaned, placed on the PDMS molds, and processed by drying in a vacuum oven at 125°C for 2 h, followed by baking at 190°C for 1 h to induce polymer reflow and generate U-shaped microwells with a depth of approx. 250 µm. The resulting PC masters were subsequently replicated in PDMS using standard soft lithography procedures.

The middle layer was fabricated using dry photoresist masters as previously described.^20^ Here, SUEX films with a thickness of 75 µm were laminated onto 3 in silicon wafers and photolithographically patterned. PDMS mixture was prepared, degassed, cast onto the master molds, and cured at 80°C for 2 h. Inlet and outlet ports were created using a 1.5 mm biopsy punch (TedPella Inc., Redding, CA, USA). The top reservoir layer was formed by casting a 5mm thick PDMS slab, followed by the formation of reservoirs using a 4 mm biopsy punch.

All PDMS layers were cleaned with adhesive tape to remove particulates. The bottom and middle layers were aligned under a stereomicroscope, such that the U-shaped microwells were registered with the corresponding culture chambers. Bonding was done using oxygen plasma (20% O_2_, 50 W) for 1 min (PE-50, Plasma Etch Inc., Carson City, NV, USA). The top reservoir layer was subsequently treated using the same plasma protocol and aligned manually with the inlet and outlet ports to complete device assembly. Devices fabricated across independent batches exhibited consistent feature dimensions and trapping performance, and no qualitative differences in spheroid loading efficiency or on-chip viability were observed between batches.

### 2.3. Cell culture and device seeding

A549, H1975, and H358 cells were cultured in RPMI 1640 medium supplemented with 10% FBS and 1% 100× antibiotic-antimycotic solution. WI-38 fibroblasts were cultured in DMEM supplemented with 10% FBS and 1% 100× antibiotic-antimycotic solution. HUVECs were cultured in EGM-2 medium (EBM-2 supplemented with the complete Lonza growth factor kit). All cell types were maintained in T-flasks at 37°C in an incubator with 5% CO_2_.

Spheroids were pre-formed using the agarose microarrays as previously described ^21^. Briefly, agarose microwells (200 μm diameter, 75 μm depth) were fabricated by casting 400 µL of molten agarose into each well of a standard 24-well plate, followed by the placement of PDMS molds containing micropost arrays (negative replicas of the microwell geometry) replicated from 3D-printed masters. After 10 min, the agarose solidified, and the PDMS molds were easily removed, leaving behind a patterned microwell array. Approximately 400 microwells were formed per well. A 1 mL suspension of A549, H1975, or H358 cells at 1×10^5^ cells/mL was added to each well. Cell suspensions were mixed thoroughly immediately prior to seeding to ensure uniform cell distribution and minimize aggregation. Spheroids were cultured in RPMI 1640 medium supplemented with 10% FBS and 1% antibiotic-antimycotic solution, and formation was observed after overnight incubation in 37 °C and 5% CO_2_.

For microfluidic device seeding, either single-cell or spheroid suspensions were introduced into the device inlets using 200 µL pipette tips. To achieve ∼100 cells or one spheroid per microwell, 50 µL of single-cell suspension (50,000 cells/mL) or 200 µL of spheroid suspension (∼100 spheroids/mL) was dispensed into the side-channel ports, while the two central outlet reservoirs were left empty. For dual-population seeding, opposing inlets were used to introduce a second spheroid type. Cell and spheroid entry into the device was driven by a hydrostatic pressure gradient between the filled inlets and empty outlets, enabling capture within individual microwells. Residual cells or spheroids remaining in the channels were flushed by adding fresh medium to the inlets to reinforce the pressure gradient. Following seeding, cells and spheroids were allowed to settle into the U-shaped microwells during overnight incubation.

Culture medium was exchanged every 48 h by a passive hydrostatic exchange protocol. Reservoirs on both the inlet and outlet sides were first emptied, after which fresh medium was added to the inlet reservoirs while the outlet reservoirs were left empty, creating a pressure differential that drove slow flow through the device. Successful perfusion was confirmed by a visible rise in liquid level in the outlet reservoirs after approximately 5 min. The outlet reservoir was then emptied again, and the inlet refill-wait cycle was repeated until reservoirs on both sides contained fresh medium, resulting in a complete medium exchange. This protocol was used for all on-chip operations, including drug treatment and fluorescent staining.

### 2.4. Long-term on-chip culture and viability assessment

Spheroid viability was assessed over a 4-week culture period. A staining solution containing 3 μg/mL Calcein AM (live-cell marker), 3 μg/mL propidium iodide (PI, dead-cell marker), and 3 μg/mL Hoechst 33342 (nuclear stain) was prepared in culture medium and introduced into the device using the medium exchange protocol described above. The staining solution was incubated for 30 min at 37°C and 5% CO_2_. Live-dead fluorescence imaging was performed using a 10× objective (NA = 0.3) on an Olympus IX-83 inverted fluorescence microscope with a 16-bit sCMOS camera (Zyla 5.5, Andor Technology Ltd, Belfast, UK). Image analysis was conducted using Fiji (ImageJ) software. Spheroid viability was quantified as the ratio of Calcein AM-positive area to the combined area of Calcein AM and PI-positive regions. Spheroid size was determined from the brightfield images by measuring the projected spheroid area. Each spheroid was treated as an independent data point for calculation of the mean and standard deviation (SD). The coefficient of variation (CV) was calculated as the SD divided by the mean and expressed as a percentage. All quantitative data were analyzed and plotted using OriginPro (OriginLab).

### 2.5. On-chip drug response studies of NSCLC cell line spheroids

Pre-formed A549, H358, and H1975 spheroids were loaded into the microfluidic devices as described earlier and cultured on-chip under standard conditions for 48 h to allow equilibration within the U-shaped microwells. Following equilibration, devices were exposed to targeted inhibitors for 72 h under two media conditions: standard culture medium and a conditioned medium. For KRAS G12C inhibition, H358 and A549 spheroids were treated with the KRAS G12C inhibitor adagrasib. For EGFR inhibition, H1975 and A549 spheroids were treated with the EGFR inhibitor osimertinib. In all cases, inhibitors were administered at final concentrations of 0, 250, 500, and 1000 nM.

Conditioned medium was prepared as a 1:1 (v/v) mixture of RPMI 1640 and either WI-38 or HUVEC -conditioned medium, as indicated for each experiment. Drug treatment was performed using the passive hydrostatic exchange protocol described above. Cytotoxicity was assessed using the live-dead staining protocol described previously. Spheroid viability was quantified as relative viability, defined as the ratio of the mean viability of spheroids in each treated group to that of the corresponding untreated control group, expressed as a percentage. All spheroids within each device were included in the analysis. Data were processed and visualized using Origin Pro (OriginLab).

### 2.6. Controlled continuous perfusion in microfluidic chip

Continuous perfusion within the microfluidic device was established using a programmable syringe pump connected via flexible tubing. Tygon tubing 0.020” id × 0.060” od (VWR) was used to connect the device inlets to the pump. A Legato 200 syringe pump (KD Scientific) was used to deliver culture medium or drug-containing solutions through the tubing. For perfusion experiments, one inlet on a side channel was connected to a syringe mounted on the pump, while the outlet was defined as the open port on the opposing side channel. All remaining ports were sealed using short segments of tubing secured with acrylic adhesive to prevent leakage and to maintain unidirectional flow across the microwell array. The flow rate was set to 70 nL/min for both culture medium and drug solution perfusion. This flow rate was selected to generate an estimated wall shear stress of approximately 4.16 × 10^-3^ dyn/cm^2^, calculated based on channel dimensions and volumetric flow rate, and was within the range of shear stresses previously reported for microphysiological tissue culture models.^22,23^ Perfusion was maintained continuously for the duration specified in each experiment.

### 2.7. On-chip culture and drug response evaluation of PDOs

PDOs F231 and F671, obtained from NCI Patient-Derived Models Repository (PDMR), were expanded off-chip in 6A Type medium. For initial expansion, organoids were suspended in Basement Membrane Extract Type 2 (BME2) and seeded as domes, following repository-recommended protocols. Cultures were maintained in 37°C in incubator with 5% CO_2_. For on-chip experiments, intact PDOs were released from BME2 and loaded into the microfluidic devices using the same trapping and seeding protocol described for spheroids. Following loading, PDOs were maintained under static culture conditions using the previously described passive hydrostatic medium exchange protocol. PDO viability and size were assessed immediately after loading (day 0) and at weeks 1, 2, and 3 post-seeding. Live-dead fluorescence staining was performed using Calcein AM, PI, and Hoechst 33342 at the same concentrations described for NSCLC spheroids. Imaging and image analysis were conducted using the same instrumentation and workflow described above. PDO size was quantified from projected area measurements, and viability was calculated from fluorescence images.

For drug response studies, F231 and F671 PDOs were allowed to equilibrate on-chip for 24 h prior to treatment. Organoids were then treated with the KRAS G12C inhibitor adagrasib for 72 h. Adagrasib was administered at final concentrations of 0, 250, 500, 1000, and 2000 nM under two media conditions: standard culture medium and conditioned medium composed of a 1:1 (v/v) mixture of RPMI 1640 and WI-38-conditioned medium. Drug treatment was performed using the passive hydrostatic exchange protocol described earlier.

Cytotoxicity was assessed using the live-dead fluorescence staining. PDO viability was quantified as relative viability, defined as the ratio of the mean viability of organoids in each treatment group to that of the corresponding untreated control group. Statistical comparisons between treatment conditions were performed using two-way ANOVA to evaluate the effects of adagrasib concentration and media composition. All quantitative data were analyzed and visualized using OriginPro (OriginLab).

## 3. Results

### 3.1. Design and operation of the microfluidic NSCLC MPS

The microfluidic MPS was designed to support parallel culture of tumor spheroids and PDOs, controlled drug delivery, and modular incorporation of TME components within a single platform. The device architecture was intentionally configured to support multiple modes of spheroid generation and loading, uniform size control, spatial compartmentalization, and scalable drug testing. Together, these features provide a consistent and well-controlled baseline for interrogation of microenvironment-driven drug resistance.

The system consisted of a PDMS device with 5 parallel culture units for 3D tumor culture and drug testing. Each unit consisted of a bottom layer patterned with 30 U-shaped microwells and an aligned top layer containing 30 culture chambers connected to a central main channel through arrays of microfabricated posts (**Figure 1**). The microwells had a diameter of approximately 400 µm and a depth of 250 µm, dimensions selected to promote formation and retention of compact spheroids and organoids while limiting vertical displacement during perfusion or media exchange. Two lateral side channels with microfabricated filters connected to each culture unit to enable spheroid loading and controlled media delivery (**Figure 2**).

**Figure 1.**
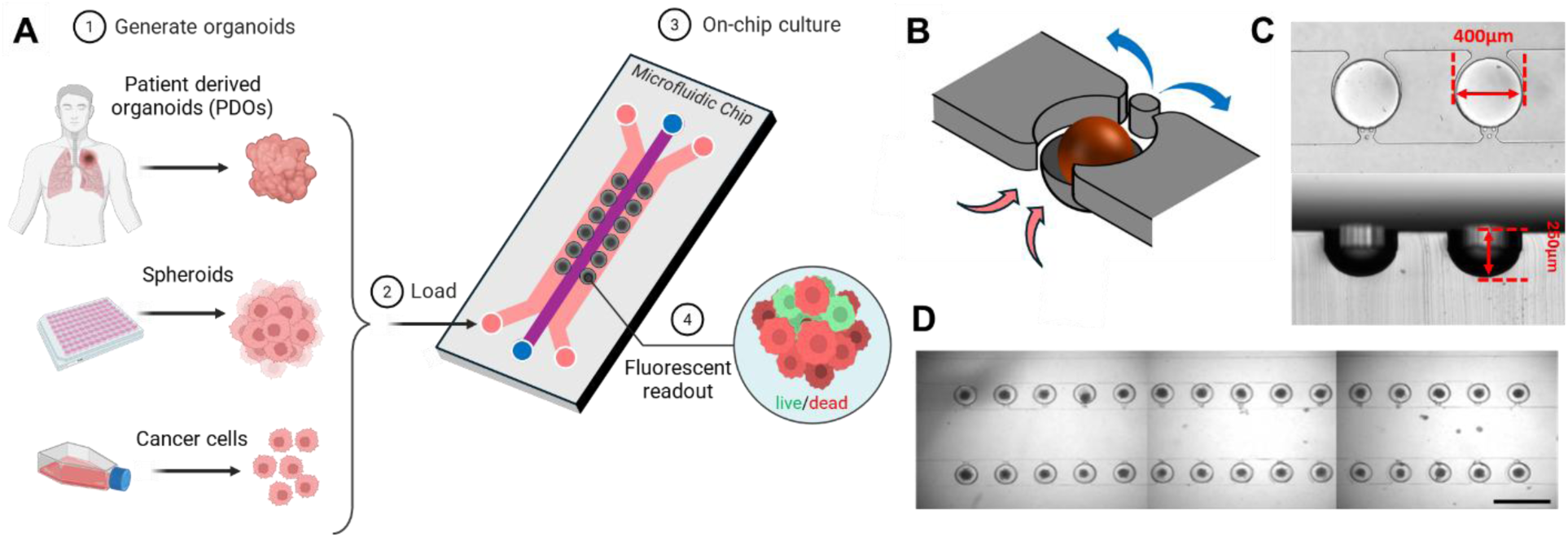
Microphysiological system concept and single-chamber operating principles. The microphysiological platform is designed to enable controlled tumor size, modular integration of tumor microenvironment components, and quantitative assessment of therapeutic response. **(A)** Conceptual workflow of the platform. In Step 1, 3D tumor models are generated from patient-derived organoids, spheroids preformed in U-well microwell arrays, or aggregates formed directly from cancer cell cultures. In Step 2, spheroids or organoids are loaded into the microfluidic device. In Step 3, on-chip culture and drug treatment are performed under static or perfused conditions. In Step 4, therapeutic response is assessed using fluorescence-based viability or immunofluorescence assays, and conditioned media can be collected through the central channel for downstream analysis of soluble factors. **(B)** Isometric 3D schematic of a single culture chamber illustrating local trapping and perfusion mechanics. The organoid is partially seated within a U-shaped microwell, with the upper portion extending into the culture chamber. A microfabricated post prevents organoid displacement under flow, while red and blue arrows indicate media inflow and outflow, respectively. **(C)** Representative top-view and cross-sectional images of U-shaped microwells and post arrays, highlighting physical confinement used to control spheroid positioning and geometry. Culture chambers have a diameter of 400 µm and microwells have a depth of 250 µm. **(D)** Stitched brightfield image of a fabricated device showing the central media channel and uniformly populated culture chambers, demonstrating consistent loading and spatial organization of spheroids or organoids.

**Figure 2.**
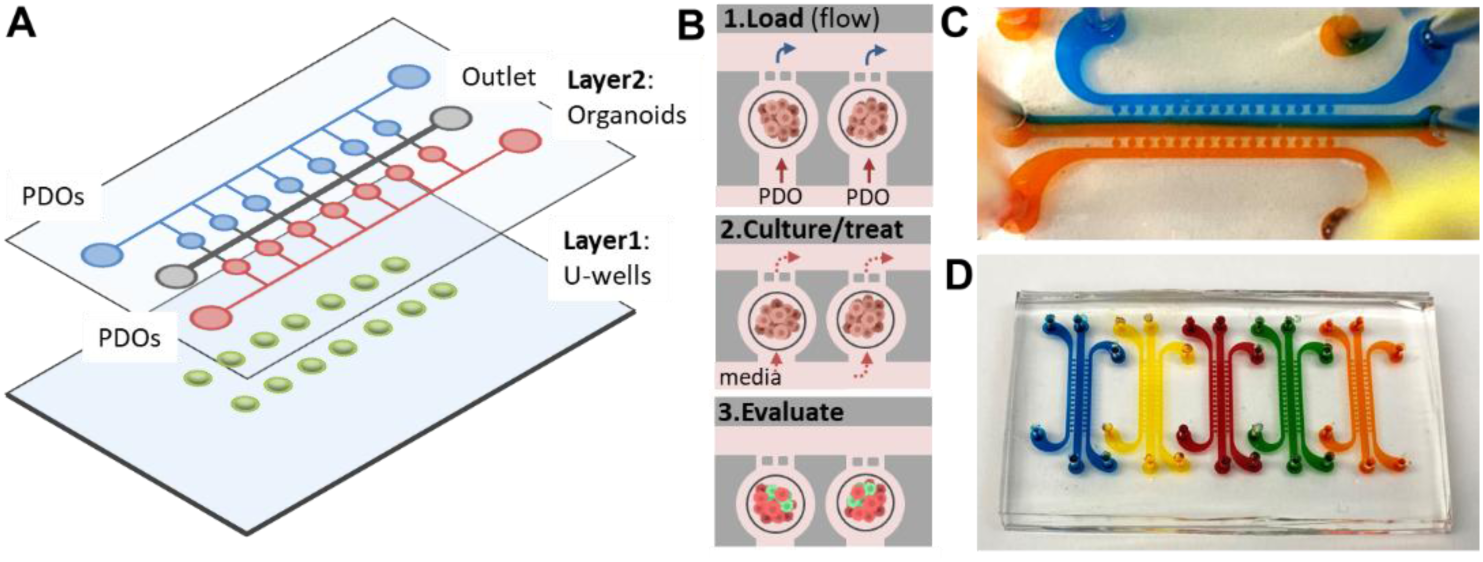
Architecture and operation of the MPS. The platform architecture supported independent channel control and parallelized drug exposure across multiple culture units, enabling scalable, internally controlled drug testing. **(A)** Exploded schematic of the full device showing the bottom microwell layer and the aligned top culture chamber layer. Lateral microchannels enable loading of spheroids or patient-derived organoids, allowing identical or distinct tissue types to be introduced through each channel. The central channel enables the collection of conditioned media for the analysis of soluble factors and can subsequently be repurposed to culture tumor microenvironment cell types for studies of *de novo* resistance. **(B)** Top-down schematic illustrating array-level spheroid and organoid loading and hydrodynamic trapping across adjacent culture chambers. U-shaped microwells capture individual spheroids or organoids, while post arrays guide loading and prevent displacement during flow. This configuration supports tumor monocultures or tangled co-cultures and enables shared media exchange, optional perfusion, and paracrine communication across chambers. **(C)** Photograph of a fabricated device visualized using colored dyes to illustrate independent media delivery to separate channels, alongside a brightfield image of a loaded device showing uniform occupancy of microwells by spheroids. **(D)** Photograph of a fabricated chip containing five parallel MPS devices, each visualized with a distinct colored dye, enabling simultaneous assessment of a four-point drug concentration series plus control within a single experiment.

Pre-formed spheroids or organoids introduced through the side channels were efficiently captured within individual microwells by a combination of physical confinement by the posts and hydrodynamic resistance (**Figure 1**). Once a microwell was occupied, the local increase in flow resistance diverted subsequent spheroids toward unoccupied wells, promoting uniform loading across the array. To further minimize positional bias and prevent preferential loading near the inlets, a dual-sided loading strategy was implemented in which spheroid suspensions were introduced sequentially from both ends of the side channels. Using this approach, the coefficient of variation (CV) of spheroid size within the device was approximately 23.4 ± 3.7 %, establishing a level of uniformity suitable for quantitative comparison of drug responses across conditions.

At the device level, the parallel layout enabled uniform culture conditions and independent control of experimental parameters across multiple culture units (**Figure 2**). The device’s parallel layout enabled simultaneous delivery of up to four distinct drug concentrations to different culture units by independently controlling inlet flows. This architecture supported both static culture and controlled perfusion, allowing interrogation of drug response under transport conditions that more closely approximate *in vivo* exposure. In selected experiments, a common drug mixing channel was incorporated upstream of the culture units to generate well-defined concentration gradients across the array, further expanding the range of pharmacological perturbations that could be applied within a single device.

To ensure precise control over initial spheroid size and maturity, and to facilitate direct comparison between open and microfluidic culture formats, the microfluidic platform was integrated with an agarose microwell array used for off-chip spheroid formation. H1975 and A549 spheroids generated in agarose exhibited high viability across a range of seeding densities, confirming the robustness of this approach for producing standardized starting populations.

Seeding at 20,000 or 50,000 cells/ml yielded predictable differences in spheroid size, with both cell lines maintaining viability >80% after one week and > 76% after two weeks under all conditions.

Spheroid growth kinetics in agarose were strongly dependent on initial seeding density. H1975 spheroids increased in area by a factor of ∼4 between the first and second weeks in culture, while A549 spheroids exhibited 2-3 fold increases depending on the starting concentration. These results established agarose microwells as a tunable, high viability system for generating spheroids of defined size prior to microfluidic loading. After initial culture in agarose, spheroids were transferred into the microfluidic device and trapped within U-shaped microwells using the dual-sided loading protocol without observable disruption of spheroid integrity.

This integrated workflow enabled systematic comparison of spheroid behavior and drug response between static and microfluidic environments, while maintaining consistent initial conditions. By explicitly decoupling spheroid generation from on-chip culture and drug delivery, the platform enabled standardized initial spheroid conditions from which the effects of TME-derived cues, including stromal and endothelial signaling, could be quantitatively assessed in subsequent experiments. From an engineering perspective, the parallel unit layout, independent channel control, and modular loading strategy together define a scalable MPS architecture that supports simultaneous testing of multiple tumor models and drug conditions within a single device, while preserving uniform hydrodynamic loading, controlled transport, and consistent exposure across experimental conditions.

### 3.2. On-chip generation, loading, and long-term maintenance of NSCLC spheroids and PDOs

Having established the design, operation, and loading performance of the microfluidic MPS, we next applied the platform to biologically relevant 3D tumor models to evaluate long-term culture, efficient on-chip trapping, and defined geometric confinement of tumor spheroids and PDOs under standardized conditions (**Figure 1A**). Rather than emphasizing spheroid formation on-chip, subsequent experiments focused on the controlled trapping and culture of pre-formed spheroids and intact PDOs, which served as the basis for all downstream drug response studies.

The microfluidic microwell array supported efficient trapping and long-term culture of pre-formed NSCLC spheroids generated from H358 and A549 cell lines. Spheroids introduced through the side channels were reproducibly captured within individual U-shaped microwells without detectable structural disruption. Longitudinal live-dead staining demonstrated that spheroids derived from both cell lines remained highly viable during on-chip culture (**Figure 3A-C**). Viability exceeded 93% after 1 week and remained >85% after 2 weeks for both lines, with A549 spheroids maintaining viability >95% even after 4 weeks. Over time, spheroids progressively compacted within the microwells. A549 spheroids exhibited a gradual decrease in projected area during the first 3 weeks, consistent with increased cellular packing, whereas H358 spheroids stabilized in size following initial aggregation during the first week.

**Figure 3.**
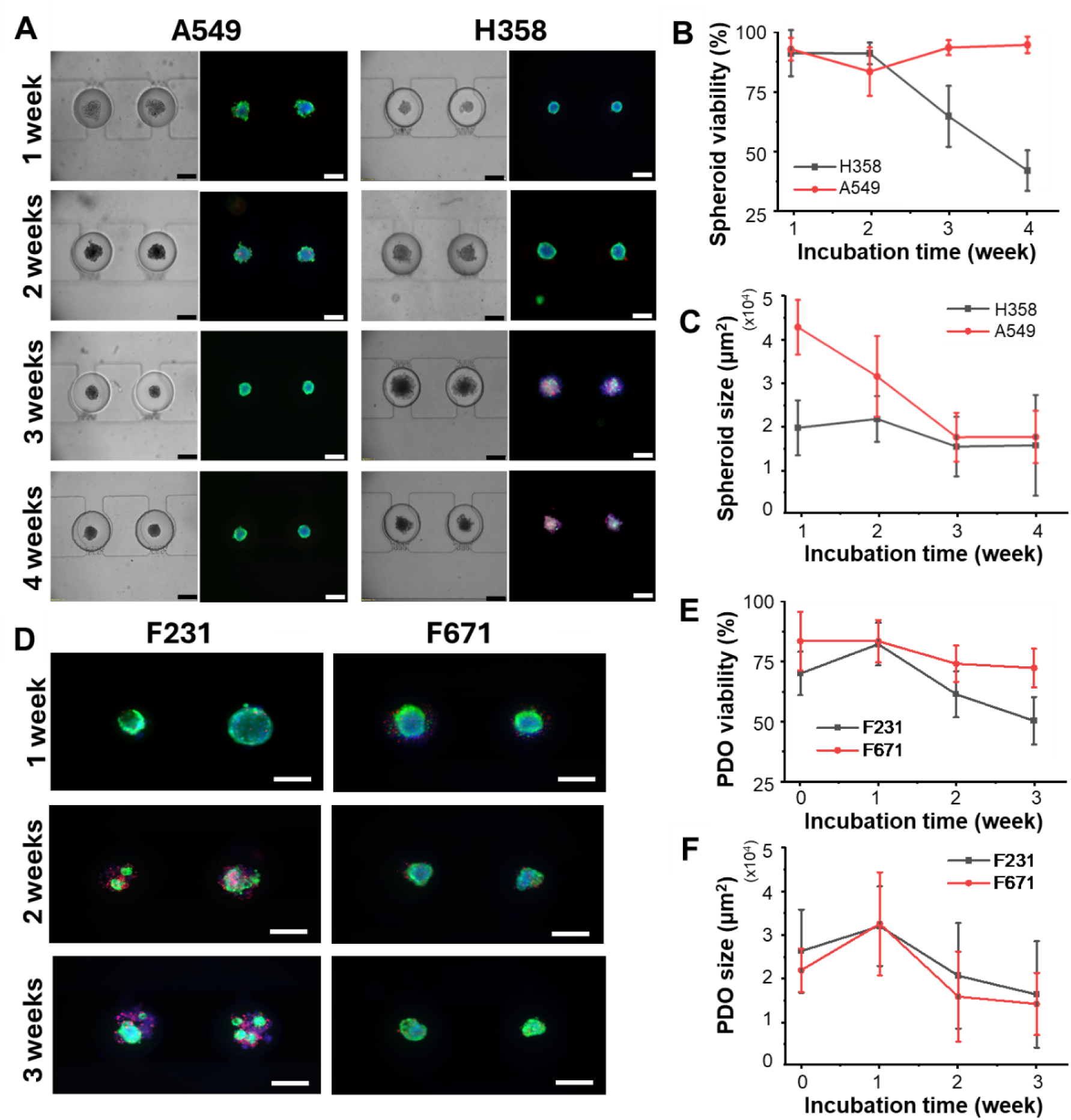
Long-term viability and morphology of NSCLC spheroids and PDOs on-chip. The MPS sustains long-term, high-viability 3D NSCLC cultures with controlled morphology, supporting quantitative drug studies. **(A)** Representative live-dead staining of A549 and H358 spheroids cultured on-chip for 1 to 4 weeks. Live cells are shown in green (Calcein AM), dead cells in red (propidium iodide), and nuclei in blue (Hoechst 33342). Scale bar, 200 µm. **(B)** Quantification of spheroid viability over time demonstrates sustained high viability during extended culture for both cell lines (*n* = 50 spheroids per condition). **(C)** Projected spheroid area over 4 weeks illustrates progressive compaction. **(D)** Representative live-dead staining of KRAS G12C mutant PDOs F231 and F671 after 1, 2, and 3 weeks of on-chip culture. Scale bar, 1 mm. **(E)** Quantification of PDO viability over time shows maintenance of high viability during the first 2 weeks, followed by a gradual decline. **(F)** Projected PDO area over time demonstrates reduced size heterogeneity and stable confinement within the microwells.

In addition to supporting long-term viability, physical confinement within the U-shaped microwells imposed consistent geometric constraints on spheroid morphology. High-resolution light-sheet imaging revealed that spheroids cultured on-chip deviated from idealized spherical geometry. Instead, aggregates adopted a constrained ellipsoidal morphology characterized by a curved basal surface conforming to the microwell and a flattened apical surface facing the culture chamber (**Figure 4**). This non-spherical geometry reflects direct mechanical interaction with the microenvironment and indicates that commonly assumed spherical symmetry may not accurately describe transport, diffusion, or drug-penetration dynamics in microwell-based tumor models. Because all subsequent drug response studies were performed using spheroids confined within the same microwell geometry, these physical constraints apply uniformly to the transport and drug-penetration conditions evaluated throughout this study.

**Figure 4.**
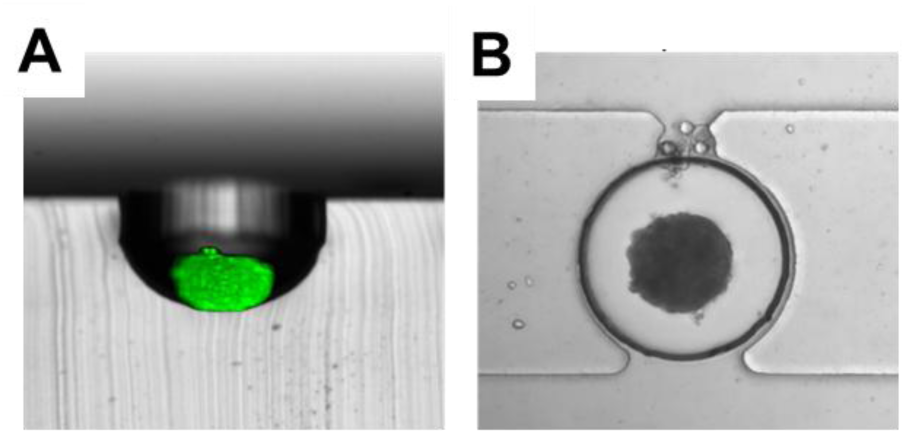
Spheroid geometry within U-shaped microwells. Physical confinement within microwells alters spheroid geometry, challenging common assumptions of spherical tumor morphology used in diffusion and drug transport models. **(A)** Composite side-view image showing a light-sheet fluorescence image of an H358 spheroid overlaid with a brightfield image of a cross-sectioned PDMS microwell. The composite illustrates both the U-shaped microwell cross-section and the resulting positioning and geometry of the confined spheroid, revealing a curved basal surface conforming to the microwell and a flattened apical surface facing the culture chamber. **(B)** Corresponding brightfield image of the same spheroid within the microwell, illustrating overall morphology and positioning within the culture chamber.

Beyond cancer cell line spheroids, the platform supported loading and culture of intact NSCLC PDOs, enabling evaluation of clinically relevant tumor material under controlled microfluidic conditions. PDOs derived from KRAS G12C-positive tumors, F231 and F671, were loaded into microwells and maintained under static conditions for up to three weeks. Live-dead staining showed that PDOs retained high viability during the first two weeks of on-chip culture, with a gradual decline thereafter (**Figure 3D-F**). F231 PDOs exhibited viability of 69.4 ± 9.1% on day zero, 81.7 ± 8.9% at week one, 60.7 ± 9.7% at week two, and 49.6 ± 9.9% at week three. F671 PDOs showed 83.1 ± 12.2% viability at day zero, 82.9 ± 8.9% at week one, 73.5 ± 7.6% at week two, and 71.7 ± 8.1% at week three.

Importantly, the microfluidic microwell architecture substantially reduced PDO size heterogeneity relative to conventional Matrigel dome cultures. Immediately after trapping, the coefficient of variation in PDO size was 36.0% for F231 and 22.6% for F671, and these distributions remained comparably narrow after one week of on-chip culture. In contrast, Matrigel dome cultures yielded markedly broader size distributions, with coefficients of variation exceeding 120% (e.g., 126.4% for F231 after 7 days, *n* = 3 ^18^). This pronounced reduction in size variability establishes a uniform and quantitative baseline for downstream drug screening, minimizing the confounding effects of initial organoid size on treatment response.

### 3.3. Static vs. perfused culture preserves baseline tumor-intrinsic drug sensitivity

Having established that spheroids experience imposes uniform physical constraints on spheroid geometry, we next evaluated whether static culture conditions were sufficient to preserve baseline tumor-intrinsic drug sensitivity or whether continuous perfusion was required to avoid transport-related artifacts. To this end, H358 spheroids were cultured on-chip under either static or continuously perfused conditions during exposure to the KRAS G12C inhibitor adagrasib. Perfusion was implemented by directing flow across spheroid-containing microwells from one inlet to the opposing outlet (**Figure 5A**). A constant flow rate of approximately 70 nL/min was maintained throughout the treatment period, corresponding to a low-shear stress regime of approx. 0.05 dyne/cm^2^, commonly used in microphysiological tumor culture systems.^22–24^

**Figure 5.**
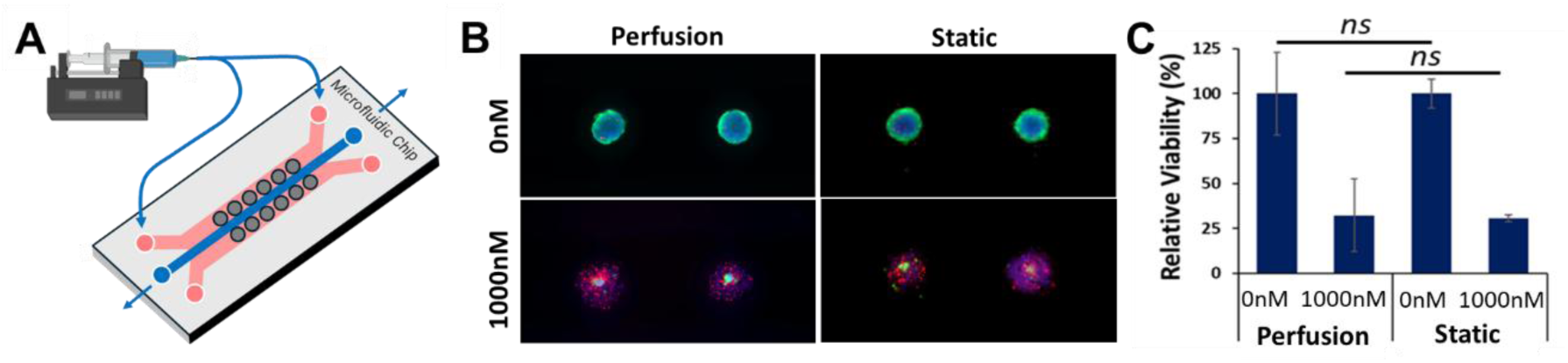
Static and perfused cultures preserve baseline KRAS G12C response in the microfluidic MPS. **(A)** Schematic illustration of the perfusion configuration used to direct flow across spheroid-containing microwells. **(B)** Representative live-dead fluorescence images of H358 spheroids cultured under perfused or static conditions during adagrasib treatment. Live cells are shown in green (Calcein AM) and dead cells in red (propidium iodide). **(C)** Quantification of spheroid viability following 72 h of adagrasib exposure under perfused and static conditions shows comparable dose-dependent responses, with no statistically significant differences between culture modes. Data are shown as mean ± SD, with each spheroid treated as an independent data point.

Comparison of spheroid viability after 72 h of adagrasib exposure revealed no statistically significant differences between static and perfused cultures across the tested concentration range (**Figure 5C**). Dose-dependent loss of viability was preserved under both conditions, and the relative sensitivity of H358 spheroids to KRAS G12C inhibition was not altered by the presence of continuous flow. These results indicate that, under the confined microwell geometry and low-shear perfusion regime used here, static culture does not confound intrinsic pharmacological response. Based on these findings, subsequent drug response experiments were conducted under static culture conditions, minimizing experimental complexity while preserving intrinsic drug sensitivity.

### 3.4. Baseline targeted therapy responses in microfluidic MPS

To define baseline tumor-intrinsic drug responses in the absence of explicit microenvironmental modulation, we first evaluated the sensitivity of H1975 and A549 spheroids to the EGFR inhibitor osimertinib in agarose microwell arrays (**Figure 6**). Spheroids were exposed to a concentration range of 125 to 2000 nM, centered around a clinically relevant level near 500 nM. Consistent with known genotypes, H1975 spheroids displayed a pronounced dose-dependent loss of viability, while A549 spheroids remained largely insensitive across all concentrations tested. At doses of 500 nM and above, H1975 spheroid viability decreased sharply, with only ∼23% viability remaining at 2000 nM, while A549 spheroids maintained viability >83% across all doses.

**Figure 6.**
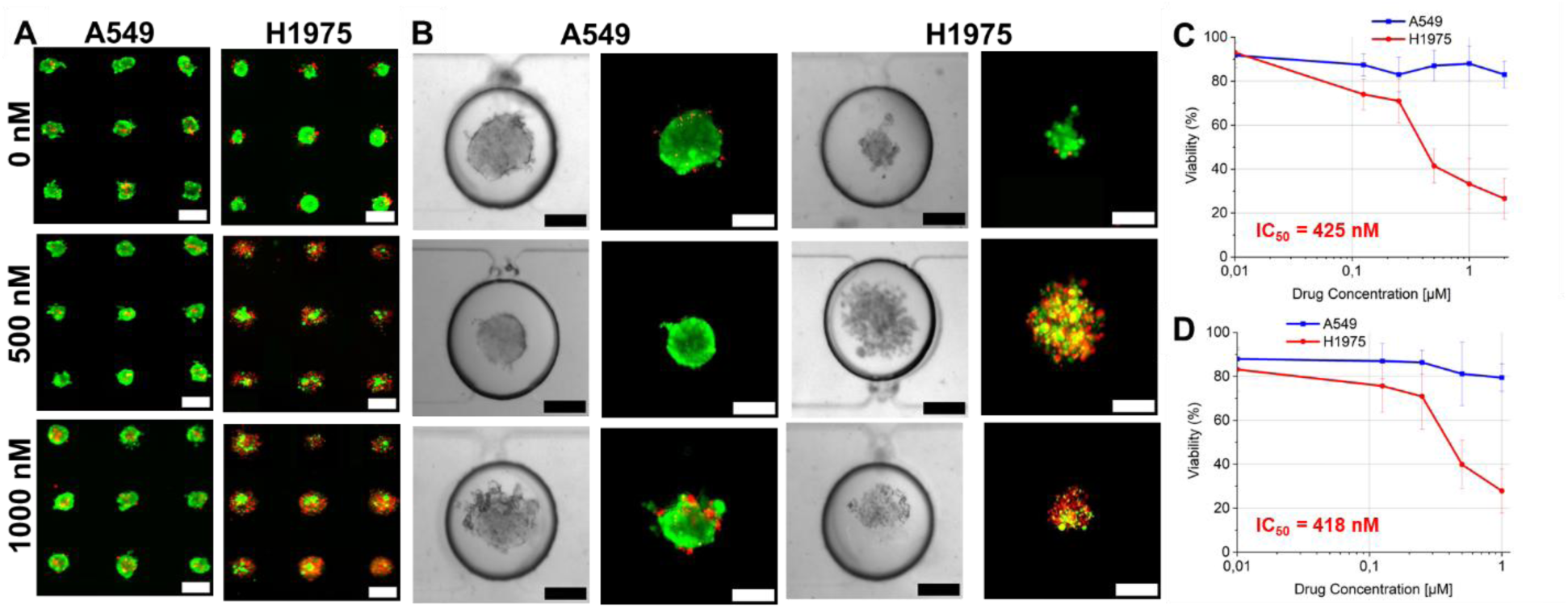
Preservation of baseline tumor-intrinsic responses to EGFR inhibition in MPS. Genotype-dependent sensitivity to targeted therapy is preserved on-chip, establishing a validated baseline prior to interrogation of tumor microenvironment effects. **(A)** Representative live/dead staining images of A549 (EGFR wild-type) and H1975 (EGFR mutant) spheroids cultured in agarose U-shaped microwells and treated with increasing concentrations of osimertinib (0-2000 nM). Live cells are shown in green (Calcein AM) and dead cells in red (propidium iodide). Scale bar, 200 µm. **(B)** Representative brightfield and live/dead fluorescence images of A549 and H1975 spheroids cultured on-chip following osimertinib treatment (0-1000 nM; concentrations selected to bracket the IC50), illustrating comparable genotype-dependent responses in the MPS. Scale bar, 150 µm. **(C)** Quantitative dose-response curves for agarose microwell cultures demonstrate no response to osimertinib in A549 spheroids and the expected dose-dependent cytotoxicity in H1975 spheroids, with an IC50 = 425 nM. **(D)** Corresponding dose-response curves for on-chip cultures show preserved drug sensitivity profiles, with A549 spheroids remaining insensitive and H1975 spheroids exhibiting a comparable IC50 = 418 nM.

Live-dead imaging revealed extensive propidium iodide uptake and disaggregation of H1975 spheroids at higher osimertinib concentrations, whereas A549 spheroids maintained tight and cohesive morphology (**Figure 6A**). Quantitative analysis of spheroid area supported these trends. Osimertinib treatment did not significantly affect A549 spheroid size, whereas H1975 spheroid area decreased to approximately 5000 µm^2^ at concentrations above 500 nM. From these data, the IC_50_ for H1975 spheroids cultured in agarose microwells was ∼425 nM, consistent with expected sensitivity to EGFR inhibition (**Figure 6C**).

We next examined whether these intrinsic drug response profiles were preserved after spheroid transfer into the microfluidic MPS. H1975 and A549 spheroids were cultured for 48 h in agarose, transferred into the microfluidic microwells, allowed to equilibrate for an additional 48 h, and then exposed to 125-1000 nM osimertinib for 72 h. As observed in agarose, A549 spheroids cultured in the MPS showed no significant cytotoxicity, maintaining compact morphology and high viability even at the highest drug concentration tested. In contrast, H1975 spheroids exhibited a clear dose-dependent loss of viability, with substantial integrity loss at 500 nM and near-complete structural disruption at 1000 nM.

The IC_50_ for H1975 spheroids cultured in the MPS was ∼418 nM, closely matching the value obtained in agarose microwells (**Figure 6D**). High-resolution imaging further revealed fragmentation and dispersal of H1975 spheroids at higher drug concentrations without the overexposure artifacts sometimes observed in well plates, underscoring the imaging advantages of the microfluidic geometry. Direct comparison of dose response curves confirmed strong agreement between agarose and the microfluidic platform in both viability and size metrics.

Although untreated H1975 spheroids exhibited slightly lower baseline viability in the MPS than in agarose (∼83.3% vs. 93.3%), this modest difference likely reflected mechanical perturbation during transfer rather than fundamental alterations in cellular sensitivity. Together, these results demonstrate that baseline tumor-intrinsic responses to EGFR-targeted therapy are preserved across agarose and microfluidic culture formats, establishing a validated reference framework for subsequent isolation of TME-mediated effects.

### 3.5. Fibroblast-derived paracrine signaling attenuates KRAS G12C inhibitor efficacy

With baseline tumor-intrinsic drug responses established and shown to be preserved within the microfluidic MPS, we next asked whether fibroblast-derived paracrine signals were sufficient to modulate KRAS G12C targeted therapy response, and whether this effect generalized from cancer cell line spheroids to PDOs. To address this question, adagrasib responses were first evaluated under standard culture conditions and then compared with responses obtained in the presence of fibroblast-conditioned medium, while maintaining identical spheroid handling, trapping geometry, and dosing protocols within the MPS workflow (**Figure 7**).

**Figure 7.**
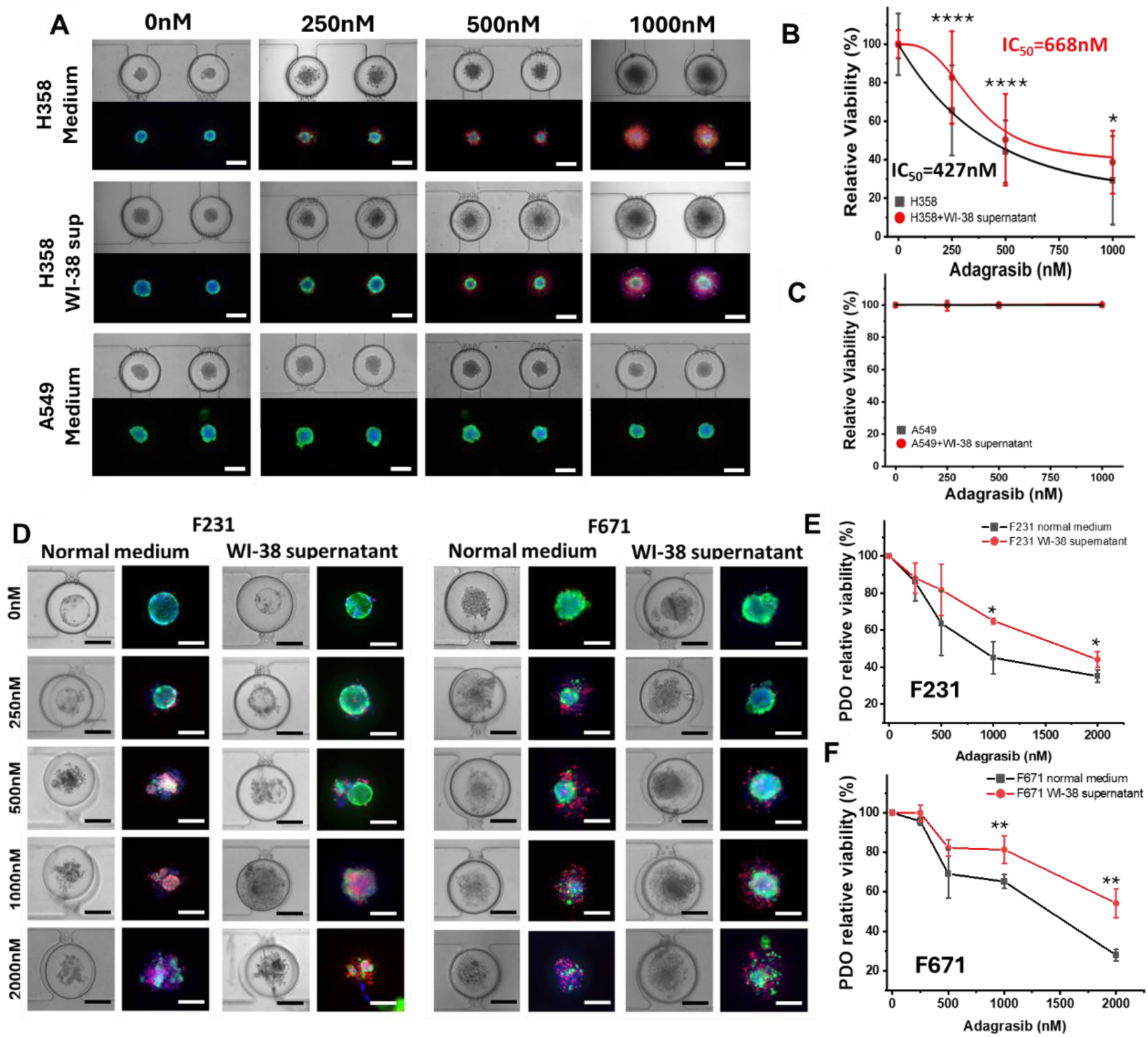
Fibroblast-derived paracrine signaling attenuates KRAS G12C inhibitor efficacy across NSCLC spheroids and PDOs. Soluble factors secreted by fibroblasts are sufficient to induce resistance to KRAS G12C inhibition in both cell line-derived spheroids and PDOs. **(A)** Representative live and dead staining images of H358 and A549 spheroids cultured in standard medium or WI-38 fibroblast-conditioned supernatant following 72 h treatment with the KRAS G12C inhibitor adagrasib. Live cells are shown in green (Calcein AM), dead cells in red (propidium iodide), and nuclei in blue (Hoechst 33342). Scale bar, 200 µm. **(B)** Quantification of H358 spheroid viability demonstrates a dose-dependent cytotoxic response to adagrasib that is significantly attenuated in the presence of WI-38 conditioned supernatant, indicating fibroblast-mediated drug resistance (*n* = 50 spheroids). **(C)** A549 spheroids, which lack the KRAS G12C mutation, show minimal response to adagrasib under both standard and conditioned media conditions, serving as a genotype-matched negative control. Statistical significance was assessed using a two-sample t-test (*p < 0.05, ****p < 0.0001). **(D)** Representative live and dead staining images of KRAS G12C mutant PODs F231 and F671 cultured in standard medium or WI-38 conditioned supernatant during adagrasib treatment. Scale bar, 200 µm. **(E)** Quantification of PDO viability reveals reduced sensitivity to adagrasib in the presence of fibroblast-conditioned supernatant for both F231 and F671. **(F)** Corresponding changes in PDO size further support fibroblast-mediated attenuation of drug response (*n* = 3 independent experiments).

We first characterized adagrasib response in H358 and A549 spheroids cultured in the MPS under standard medium conditions. H358 spheroids, which harbor a KRAS G12C mutation, exhibited a dose-dependent decrease in viability with adagrasib exposure, whereas A549 spheroids, which lack this mutation, showed minimal response across the same concentration range. This genotype-dependent behavior is consistent with KRAS G12C specificity and mirrors trends previously observed in agarose microwell cultures. Within the MPS, these paired responses served as an internal pharmacogenomic control, enabling subsequent changes in drug efficacy to be attributed to microenvironmental modulation rather than to differences in baseline sensitivity introduced by the culture format.

To test whether fibroblast-derived paracrine cues modulated KRAS inhibition in a microfluidic platform designed to support controlled microenvironmental perturbations using limited tumor material, H358 spheroids were treated with adagrasib in the presence or absence of WI-38 fibroblast-conditioned supernatant (**Figure 7A-B**). In standard media, adagrasib reduced spheroid viability in a concentration-dependent manner. In contrast, exposure to fibroblast-conditioned medium consistently attenuated cytotoxicity across the same dose range, indicating a protective effect mediated by soluble factors. Because spheroid loading, confinement geometry, drug delivery, and imaging workflows were identical to those used for baseline validation, the observed attenuation could be directly attributed to fibroblast-derived paracrine signaling rather than to device- or handling-related artifacts. This observation is consistent with our prior work in agarose microwells ^17,18^, in which WI-38 conditioned supernatant was shown to be enriched in hepatocyte growth factor (HGF) and to sustain proliferative signaling during KRAS G12C inhibition.

We next examined whether fibroblast-conditioned supernatant similarly altered adagrasib efficacy in KRAS G12C mutant PDOs cultured on-chip (**Figure 7D-F**). PDOs F231 and F671 exhibited dose-dependent sensitivity to adagrasib in standard media, and this response was substantially blunted in the presence of a 1:1 mixture of WI-38 supernatant and culture medium. The IC_50_ shifted from 830 nM to 1641 nM for F231 and from 1324 nM to 2979 nM for F671, indicating an approximately 2-fold increase in the drug concentration required to achieve comparable cytotoxicity. Importantly, because the microfluidic microwell architecture constrained organoid position and reduced size heterogeneity relative to Matrigel-based culture, these shifts could be interpreted against a more uniform baseline, strengthening the platform’s utility for quantitative drug screening in limited patient material. Ultimately, these results demonstrate that fibroblast-conditioned supernatant is sufficient to confer resistance to KRAS G12C inhibition in both cell line-derived spheroids and PDOs within the MPS. The consistency of these findings with our prior agarose-based studies supports a model in which fibroblast-secreted factors, including HGF, dominate the resistance phenotype, while additional structural complexity is not required to observe this effect in the present experimental context.

### 3.6. Endothelial-derived paracrine signaling attenuates EGFR inhibitor efficacy

Following identification of paracrine-dominant fibroblast-mediated resistance, we next asked whether endothelial-derived cues similarly modulated the response to EGFR inhibition, and whether resistance depended on soluble signaling or direct tumor-endothelium contact. To map increasing model complexity onto resistance mechanism, we compared 3 configurations that progressively increased physical interactions: 1) exposure to endothelial-conditioned supernatant, 2) tangled co-aggregation of tumor and endothelial cells, and 3) cross-platform comparison between agarose and the MPS (**Figure 8**). In these studies, HUVECs were used as a well-established endothelial model to demonstrate the platform’s capability.

**Figure 8.**
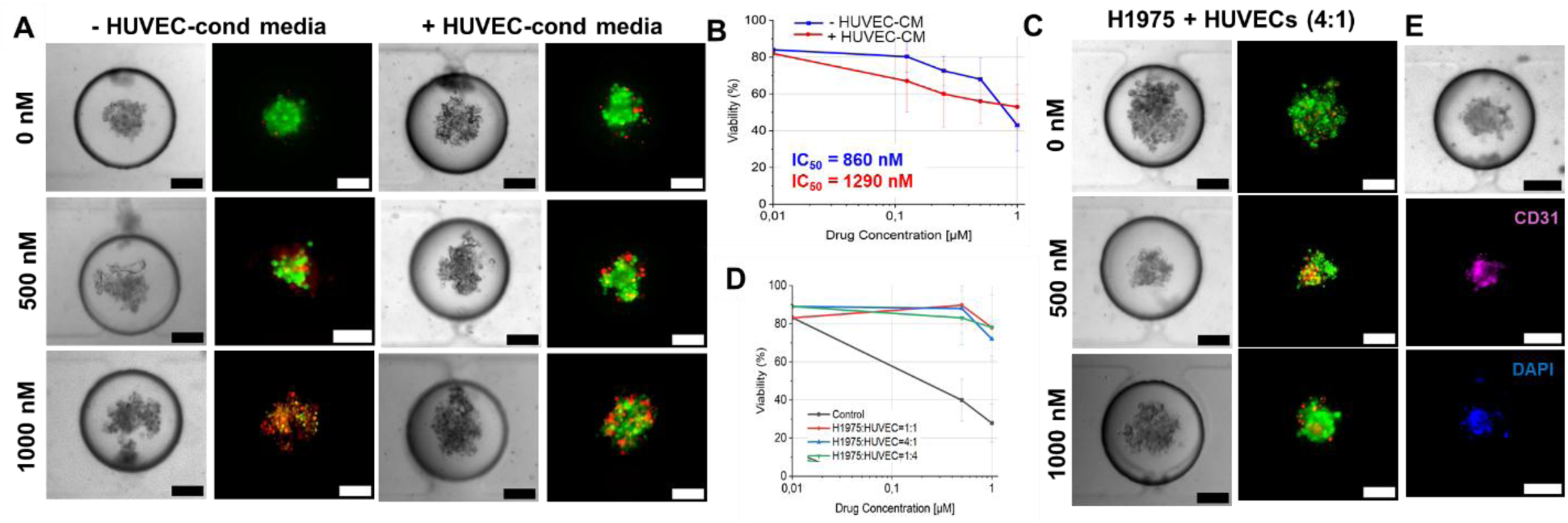
Endothelial-mediated resistance to EGFR inhibition across increasing architectural complexity. Endothelial-derived cues confer resistance to EGFR inhibition across conditioned media and co-culture configurations, indicating a dominant role for paracrine signaling, while increasing structural complexity primarily enhances morphological fidelity. (**A**) Representative brightfield and live/dead fluorescence images of H1975 spheroids cultured on-chip and treated with osimertinib (0-2000 nM) in the absence or presence of HUVEC-conditioned medium. Live cells are shown in green (Calcein AM) and dead cells in red (propidium iodide). Scale bar, 150 µm. (**B**) Quantitative dose-response curves show that exposure to HUVEC-conditioned medium reduces sensitivity to osimertinib, with the IC50 increasing from 860 nM under control conditions to 1290 nM in the presence of endothelial-derived soluble factors. (**C**) Representative images of tangled H1975-HUVEC spheroids formed at a 4:1 tumor to endothelial cell ratio and treated with osimertinib (0-1000 nM). Scale bar, 150 µm. (**D**) Dose-response analysis of tangled co-cultures demonstrates a marked attenuation of osimertinib efficacy relative to H1975 monocultures, with viability remaining >70% at the highest drug concentration tested, precluding determination of an IC50. (**E**) Immunofluorescence staining for the endothelial marker CD31 confirms incorporation of HUVEC within tangled spheroids, validating the multicellular composition of the co-culture model.

We first tested whether soluble endothelial cues altered osimertinib response in H1975 spheroids (**Figure 8A-B**). In the presence of HUVEC-conditioned medium, H1975 spheroids maintained higher viability at intermediate and high osimertinib concentrations than those treated with endothelial medium lacking conditioned factors. The IC_50_ increased from ∼860 nM under control conditions to ∼1290 nM in HUVEC-conditioned medium, indicating attenuation of EGFR inhibitor efficacy by endothelial-derived soluble signals. These results demonstrate that endothelial-secreted factors are sufficient to confer resistance to EGFR inhibition in the absence of direct cell-cell contact, paralleling observations made with fibroblast-conditioned media.

To assess whether this resistance phenotype was robust across culture formats, endothelial-conditioned media experiments were compared between agarose microwells and the microfluidic MPS. In agarose, HUVEC-conditioned medium preserved high spheroid viability across the osimertinib concentration range tested. In the microfluidic device, overall viability was modestly reduced under comparable conditions, while the qualitative resistance trend was preserved. This cross-platform concordance indicates that endothelial-mediated resistance was not an artifact of a specific culture geometry, and that soluble signaling dominated the observed response.

We next increased structural and cellular complexity by forming tangled co-cultures in which H1975 spheroids were co-aggregated with HUVECs at defined ratios prior to Osimertinib exposure (**Figure 8C-E**). Across all seeding ratios tested, tangled spheroids maintained high viability at 1000 nM osimertinib, in contrast to H1975 monocultures, which exhibited substantial drug-induced loss of viability. Notably, the protective effect did not scale with the fraction of endothelial cells incorporated into the aggregate, suggesting that resistance was triggered by qualitative exposure to endothelial-derived cues rather than by endothelial abundance.

Immunofluorescence staining for CD31 confirmed robust incorporation of endothelial cells throughout tangled spheroids, validating the structural fidelity of this co-culture configuration.

Finally, osimertinib responses in tangled co-cultures were compared directly with those observed under exposure to HUVEC-conditioned medium. Both configurations produced substantial attenuation of drug efficacy, with broadly similar preservation of viability across drug concentrations. Together, these results indicate that paracrine signaling constitutes the dominant mechanism of endothelial-mediated resistance, while increasing architectural complexity primarily enhances morphological fidelity rather than introducing a fundamentally distinct resistance pathway. Consistent with prior studies implicating growth-factor-mediated signaling in resistance to EGFR inhibition, these findings support a unifying model in which endothelial-derived soluble factors engage proliferative and survival pathways that buffer tumor cells against targeted therapy.

### 3.7. Parallelized multi-population configurations of the MPS

Having established that paracrine signaling dominates TME-mediated resistance, we next leveraged the modular architecture of the MPS to demonstrate parallelized, multi-population experimental configurations that are difficult to achieve reproducibly in conventional culture platforms. These experiments were not intended to introduce new biological variables, but rather to evaluate whether increased architectural complexity altered baseline tumor-intrinsic drug sensitivity while enabling internal controls within a single device (**Figure 9**).

**Figure 9.**
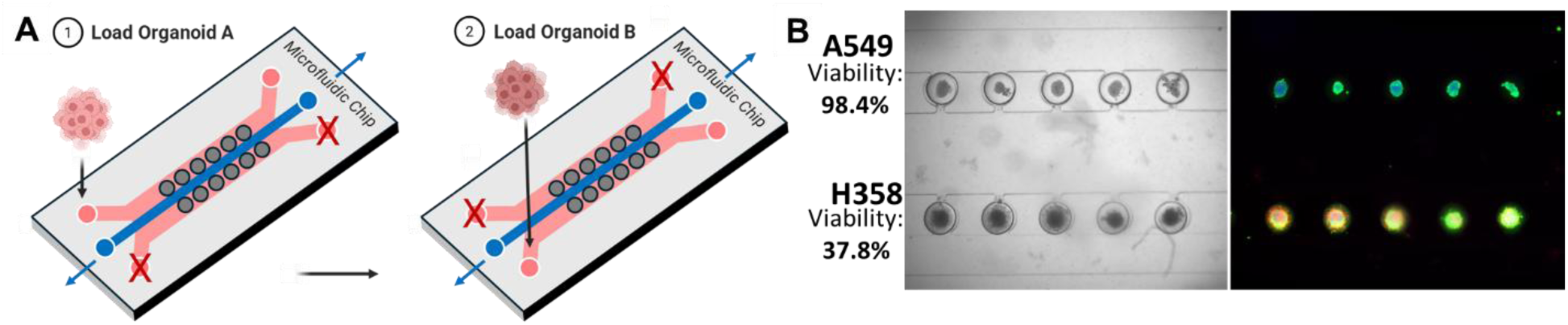
Platform-enabled parallelization and perfusion without alteration of intrinsic drug sensitivity. The MPS supports parallelized, multi-population experiments and controlled perfusion while preserving baseline tumor-intrinsic responses to targeted therapy. **(A)** Schematic illustrating dual-population loading, in which two distinct spheroid or organoid populations are subsequently introduced into separate side channels and trapped within a single device unit, enabling direct head-to-head comparison under identical culture and dosing conditions. **(B)** Representative bright field and fluorescent images of A549 and H358 spheroids following 72 h exposure to 500 nM adagrasib. A549 spheroids maintained high viability (98.4%), whereas KRAS G12C mutant H358 spheroids exhibited marked cytotoxicity (37.8% viability), consistent with genotype-dependent sensitivity. Live cells are shown in green (Calcein AM) and dead cells in red (propidium iodide).

To demonstrate spatial multiplexing, distinct NSCLC spheroid populations were loaded into opposing side channels of the same culture unit and trapped in corresponding microwells (**Figure 9A**). A549 and H358 spheroids were introduced through separate inlets and subsequently exposed to adagrasib under identical dosing conditions. As expected based on prior validation, H358 spheroids exhibited a dose-dependent decrease in viability, whereas A549 spheroids remained largely refractory (**Figure 9B**). Performing this head-to-head comparison within a single microfluidic unit minimized inter-device variability and enabled direct benchmarking of genotype-dependent drug response under identical microenvironmental and transport conditions. This configuration establishes an internal control strategy that is not readily achievable in plate-based systems and provides a foundation for future studies of inter-population crosstalk or competitive interactions.

This multi-population configuration establishes an internal control strategy that is not readily achievable in plate-based systems and provides a foundation for future studies examining inter-population variability, competitive interactions, or paracrine crosstalk under tightly controlled conditions. Together, these results demonstrate that the MPS supports parallelized comparisons of tumor populations while preserving intrinsic pharmacological behavior, reinforcing its utility as a flexible platform for systematic interrogation of tumor heterogeneity and TME modulation.

## 4. Discussion

In this work, we developed and validated a microfluidic MPS that enables controlled investigation of tumor–microenvironment interactions, with a particular focus on soluble, TME-derived cues that modulate therapeutic response in NSCLC. By first establishing a robust baseline in which cancer cell line spheroids and PDOs maintained high viability, reduced size heterogeneity, and preserved genotype-dependent responses to targeted therapies across agarose microwells and microfluidic chip, we defined a quantitative reference state against which microenvironmental effects could be assessed. Within this controlled framework, fibroblast- and endothelial-derived cues consistently attenuated responses to KRAS G12C and EGFR inhibition across multiple culture configurations, indicating a dominant role for paracrine signaling.

A key outcome of this study is the demonstration that baseline tumor-intrinsic pharmacological behavior was preserved within the microfluidic MPS. EGFR-mutant H1975 and KRAS G12C-mutant H358 spheroids exhibited dose-dependent sensitivity to osimertinib and adagrasib, respectively, whereas A549 spheroids remained refractory, consistent with their mutational status. IC_50_ values measured on-chip closely matched those obtained in agarose microwells, indicating that microwell confinement, microfluidic transport, and device materials did not intrinsically alter drug sensitivity. This concordance established a validated reference in which deviations in response could be attributed to microenvironmental modulation rather than platform-induced artifacts. Compared with conventional Matrigel-based PDO cultures,^25^ which exhibit substantial size variability and limited control over transport, this system provides a more reproducible and quantitative framework for drug response analysis.

Spheroids cultured within the U-shaped microwells exhibited non-spherical, ellipsoidal morphologies. Although this geometry was visualized in detail for H358 spheroids, similar deviations from idealized spherical structure are commonly observed in confined microscale culture systems and are not expected to be specific to a particular cell line. Variations in spheroid morphology may also reflect cell-dependent properties, including extracellular matrix deposition and intercellular cohesion. These observations highlight the importance of accounting explicitly for physical constraints when interpreting transport and drug-penetration behavior in microscale tumor models.

Evaluation of static vs. perfused culture further supported the use of simplified operating conditions for mechanistic studies. Under the confined microwell geometry and low-shear flow regime used here, static and perfused cultures yielded comparable intrinsic drug responses, consistent with diffusion-dominated treatment. While other tumor systems may exhibit greater sensitivity to perfusion, these results indicate that static culture does not confound baseline pharmacological behavior in this platform under the conditions used in this study.

Within this validated framework, fibroblast-derived paracrine signaling emerged as a potent modulator of KRAS G12C inhibitor efficacy in both cell line spheroids and PDOs, with IC_50_ values shifting approximately 2-fold in the presence of WI-38-conditioned medium, consistent with prior studies.^18^ Endothelial-derived cues similarly attenuated EGFR inhibitor response, with comparable resistance observed across conditioned media and tangled co-culture configurations. In both cases, resistance did not scale with stromal or endothelial abundance, indicating that qualitative exposure to soluble factors was sufficient to buffer tumor cells against targeted therapy. Together, these results support a unifying model in which paracrine signaling constitutes the dominant mechanism of TME-mediated resistance under the conditions examined, while increasing architectural complexity primarily enhances morphological fidelity rather than introducing distinct resistance pathways.

Conventional PDO Matrigel dome cultures preserve tumor phenotype but exhibit substantial size heterogeneity and limited control over transport, complicating quantitative analysis. Microwell systems, including our prior work with agarose microwells,^17,18,26,27^ address size uniformity and reproducibility, but lack dynamic control over perfusion and spatial organization. At the other end of the spectrum, lung-on-chip systems, vascularized organoid platforms, and immune-competent tumor-on-chip models achieve higher physiological fidelity but often require complex fabrication and culture workflows that limit scalability and mechanistic interpretation.^28^ The MPS presented here balances these trade-offs by enabling controlled, scalable interrogation of TME interactions while preserving quantitative reproducibility. In particular, the results demonstrate that key resistance phenotypes can be captured at the level of soluble signaling without requiring fully vascularized or immune-competent architectures.

Although this study highlights the dominant role of paracrine signaling in resistance to KRAS G12C and EGFR-targeted therapies, these mechanisms represent only one component of the clinically observed resistance landscape in NSCLC. In patients, resistance to KRAS G12C inhibitors commonly arises through adaptive receptor tyrosine kinase signaling, reactivation of downstream MAPK pathways, MET or EGFR pathway activation, and genomic or phenotypic heterogeneity within tumors.^29–31^ Similarly, resistance to EGFR tyrosine kinase inhibitors (TKIs) involves secondary mutations, clonal evolution, histologic transformation, and bypass signaling networks.^32,33^ Importantly, stromal and vascular components of the TME contribute to this clinical resistance by secreting growth factors, including hepatocyte growth factor and related ligands, that sustain proliferative and survival pathways despite oncogene inhibition. The ability of this MPS to isolate and quantify TME-driven paracrine resistance mechanisms in a controlled setting provides a complementary, translationally relevant framework for dissecting how these tumor-intrinsic and extrinsic factors interact.

Several limitations of the current study should be considered. The platform focuses on fibroblast- and endothelial-derived cues and does not incorporate immune components, or extracellular matrix remodeling, all of which influence tumor behavior *in vivo*. Although prior work implicates growth factor-mediated signaling, including HGF, in fibroblast-driven resistance, these mediators were not directly quantified in the present system. In addition, while the platform supports spatial compartmentalization and controlled perfusion, it does not fully recapitulate dynamic vascular remodeling or immune–tumor interactions observed in more complex organ-on-chip models.

Overall, this study establishes a framework for TME modeling in which controlled paracrine perturbations serve as efficient first-line tools for identifying dominant resistance mechanisms under defined conditions, while more complex architectures are reserved for investigations of spatial organization, transport phenomena, and tissue-level fidelity. This approach provides a scalable, mechanistically informative strategy for integrating tumor biology and microenvironmental regulation into drug response studies, with direct relevance to the development and evaluation of therapeutic strategies to overcome TME-mediated resistance in NSCLC.

## 5. Conclusions

This study establishes a modular microfluidic MPS for controlled investigation of TME interactions in NSCLC. The platform supports loading and long-term culture of cancer cell line spheroids and intact PDOs with high viability, reduced size heterogeneity, and well-defined physical confinement. Preservation of baseline tumor-intrinsic responses to targeted therapies across agarose microwells and microfluidic chip defines a validated reference state against which TME-mediated effects can be quantitatively assessed. Using this framework, fibroblast- and endothelial-derived cues consistently attenuate responses to KRAS G12C and EGFR inhibition across multiple tumor models. Conditioned media and co-culture systems of increasing architectural complexity tend to produce similar resistance-associated phenotypes, suggesting that paracrine signaling is likely a dominant factor in TME-mediated protection. Increased structural complexity improved morphological fidelity and contextual relevance, but did not fundamentally alter therapeutic response. By enabling systematic separation of baseline drug sensitivity from TME-mediated modulation under controlled geometric and transport conditions, this MPS provides a scalable and versatile platform for dissecting tumor-microenvironment interactions. More broadly, the hierarchical modeling strategy demonstrated here offers a rational framework for selecting appropriate levels of complexity in preclinical tumor models and for guiding the development of combination therapeutic strategies that target both tumor-intrinsic vulnerabilities and dominant paracrine resistance pathways in NSCLC.

## Notes

### Competing Interest Statement

The authors have declared no competing interest.

